# Multiscale Detrended Cross-Correlation Coefficient: Estimating Coupling in Nonstationary Neurophysiological Signals

**DOI:** 10.1101/2024.04.16.589689

**Authors:** Orestis Stylianou, Gianluca Susi, Martin Hoffmann, Isabel Suárez-Méndez, David López-Sanz, Michael Schirner, Petra Ritter

## Abstract

The brain consists of a vastly interconnected network of regions, the connectome. By estimating the statistical interdependence of neurophysiological time series, we can measure the functional connectivity (FC) of this connectome. Pearson’s correlation (*r*_P_) is a common metric of coupling in FC studies. Yet *r*_P_ does not account properly for the non-stationarity of the signals recorded in neuroimaging. In this study, we introduced a novel estimator of coupled dynamics termed multiscale detrended cross-correlation coefficient (MDC_3_). Firstly, we showed that MDC_3_ had higher accuracy compared to *r*_P_ using simulated time series with known coupling, as well as simulated functional magnetic resonance imaging (fMRI) signals with known underlying structural connectivity. Next, we computed functional brain networks based on empirical magnetoencephalography (MEG) and fMRI. We found that by using MDC_3_ we could construct networks of healthy populations with significantly different properties compared to *r*_P_ networks. Based on our results, we believe that MDC_3_ is a valid alternative to *r*_P_ that should be incorporated in future FC studies.

**Author Summary:** The brain consists of a vastly interconnected network of regions. To estimate the connection strength of such networks the coupling between different brain regions should be calculated. This can be achieved by using a series of statistical methods that capture the connection strength between signals originating across the brain, one of them being Pearson’s correlation (*r*_P_). Despite its benefits, *r*_P_ is not suitable for realistic estimation of brain network architecture. In this study, we introduced a novel estimator called multiscale detrended cross-correlation coefficient (MDC_3_). Firstly, we showed that MDC_3_ was more accurate than *r*_P_ using simulated signals with known connection strength, as well as simulated brain activity emerging from realistic brain simulations. Next, we constructed brain networks based on real-life brain activity, recorded using two different methodologies. We found that by using MDC_3_ we could construct networks of healthy populations with significantly different properties compared to *r*_P_ networks. Based on our results, we believe that MDC_3_ is a valid alternative to *r*_P_ that should be incorporated in future studies of brain networks.

## Introduction

Neuroscientific research has undergone a profound transformation in the last 100 years. Berger’s invention of electroencephalography (EEG) (1) made it possible to record and evaluate neural activity in a non-invasive manner. Initially, studies relied on univariate (i.e., single time series) analysis of the brain dynamics. This started to change towards the end of the 20th century with the first functional connectivity (FC) studies (2,3). This new field does not rely only on anatomical connections, it rather studies functional connections that can be created between directly or indirectly coupled neuronal populations. In more mathematical terms, the brain regions are considered nodes on a graph, interconnected by edges (4). These edges are defined by the statistical relationship of the neuronal time series under investigation.

Several different FC estimators have been introduced with Pearson’s correlation (*r*_P_) being one of the first applied in FC studies (2,3). Some drawbacks of this method (e.g., unreliable assessment of non-linear relationships) and the growing interest in exploring other aspects of FC, lead to the introduction of newer methodologies such as phase locking value (PLV) (5), phase lag index (PLI) (6), synchronization likelihood (SL) (7) and mutual information (MI) (8,9). The use of different FC estimators can greatly influence the topology of the networks (10–12). Such differences can be especially problematic when non-healthy populations are being investigated, – e.g., in Alzheimer’s disease patients (13) – complicating the reproducibility and meta-analysis of studies. It is then important that an informed choice is made before selecting an FC estimator. Nevertheless, *r*_P_ is still widely used (14) due to its simplicity and interpretability. An important advantage of *r*_P_ is the capacity to identify positive and negative correlations, which is not always the case with other estimators.

Signals can be divided into two categories: *i*) stationary and *ii*) non-stationary. A time series *X*_t_ – where *t* indicates the discrete time – is *completely stationary* when the joint probability distributions of {*X*_t1_ , *X*_t2_ , *X*_t3_ …, *X*_tn_} and {*X*_t1+k_ , *X*_t2+k_ , *X*_t3+k_ …, *X*_tn+k_} are identical for any set of time points *t*1, *t*2, *t*3…,*t*n and any integer *k*. While this definition is easily understood, it is rather unrealistic. Hence, a less strict definition for *weak stationarity* has been used to classify physiological signals. According to this, the mean and variance of a time series remain constant. In line with that, the covariance of two weakly stationary signals will also be constant throughout the propagation of time. On the other hand, non-stationary signals have varying mean and variance. Additionally, the covariance between two non-stationary signals will be time-dependent (15). **Figure 1** shows an exemplary case of these weakly-stationary and non-stationary signals. From now on, any reference to stationary signals corresponds weakly- stationary signals. Most biosignals are non-stationary (16). As a result, calculating the *r*_P_ – a standardized covariance – of two biosignals can be misleading. A solution to this issue was given with the introduction of the detrended cross-correlation coefficient (DCCC) (17). DCCC makes use of the averaged variance and covariance of smaller sections of the signals (see Section “**Multiscale Detrended Cross-Correlation Coefficient**” below). In this study, we propose an extension of DCCC termed *multiscale detrended cross-correlation coefficient* (MDC_3_). Contrary to DCCC, the output of MDC_3_ does not depend on the scale (window length) resulting in easier interpretation of the results. To show this, we compared MDC_3_ to *r*_P_ using simulated time series with: *i*) known coupling and *ii*) known causal interactions [i.e., effective connectivity (EC)]. We also demonstrated the differences between the two estimators in magnetoencephalography (MEG) and functional magnetic resonance imaging (fMRI) recordings.

**Figure 1.**
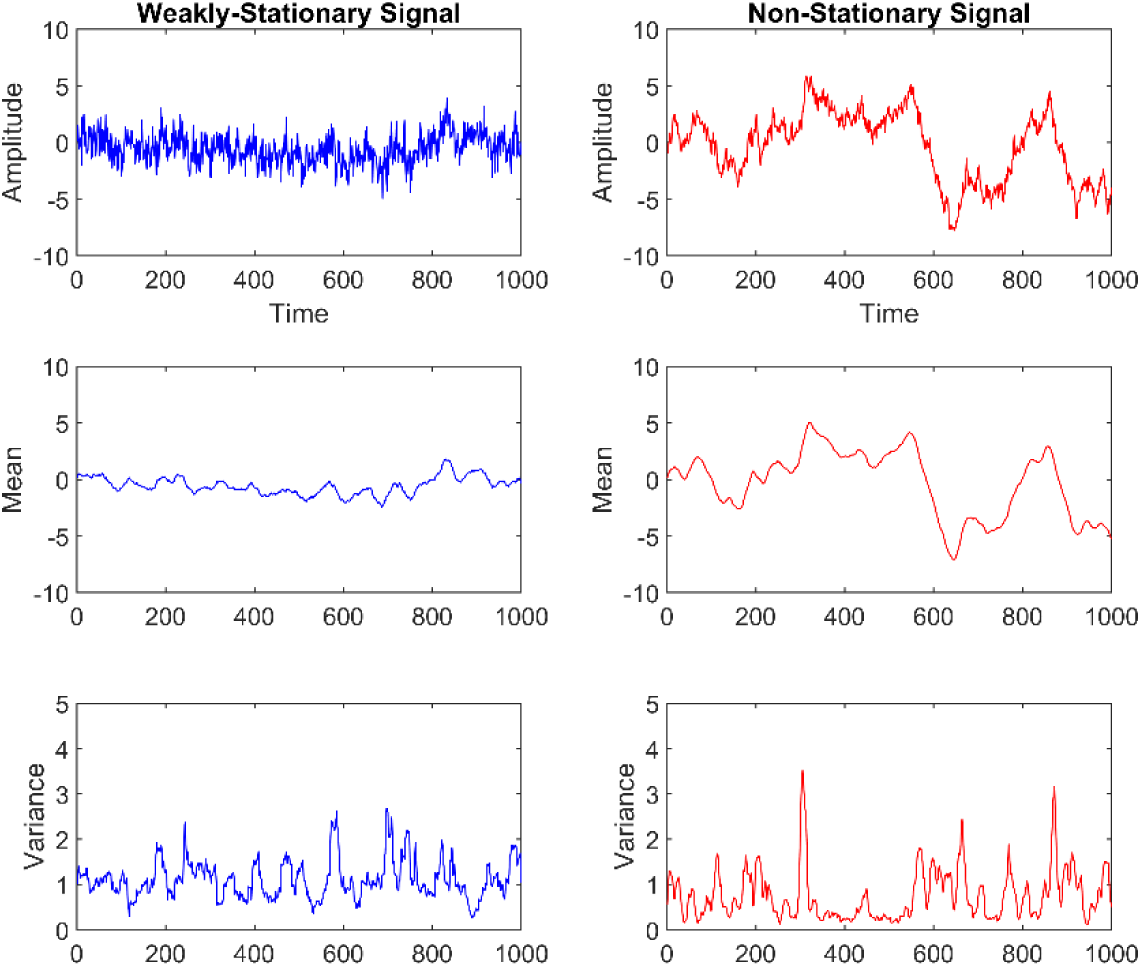
|| Example of weakly-stationary and non-stationary signals generated using auto-regressive fractionally integrated moving-average (ARFIMA) processes (see **Simulated time series**). The mean and variance of weakly-stationary signals remain constant throughout time, while they vary in non-stationary signals.

## Methods

### Multiscale Detrended Cross-Correlation Coefficient

Before introducing MDC_3_ we briefly describe DCCC (17), upon which MDC_3_ is based. DCCC was introduced as a more accurate coupling estimator between non-stationary time series. DCCC is calculated for several scales (*s*) (or window lengths) as follows. For every scale (window length), the two signals *X* and *Y* are divided into *N* non-overlapping windows of length *s*. Preliminary analysis with 50% overlapping windows did not show significant benefits compared to non-overlapping windows. For the sake of computational speed, non-overlapping windows were chosen. In every window the linear trend is removed, leaving the detrended signals 𝑋̂_𝑖_ and 𝑌̂_𝑖_, where *i* is the index of the window. Detrending is performed in order to counteract (at least partially) any spurious coupling emerging due to autocorrelation effects (18). Then, the covariance between the two signals and the variances of the two signals are estimated for every window. Finally, the ratio of average covariance and the square root of the product of average variances is calculated. **Equation 1** provides the mathematical formulation of these steps.

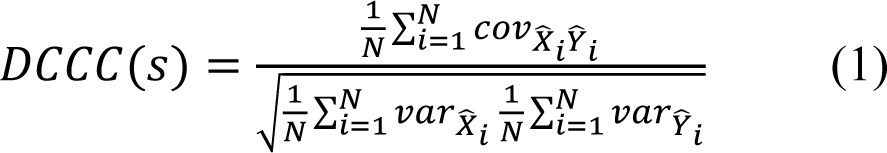

DCCC is reminiscent of *r*_P_ since both estimators range between -1 and 1 with negative values corresponding to anticorrelation and positive values corresponding to correlation (19). In 2014 Kristoufek showed that DCCC was more accurate than *r*_P_ (20) in synthetic non-stationary signals of known coupling. These results warrant the use of DCCC in FC studies, since neuronal time series are non-stationary (16). Unfortunately, the use of a multitude of scales (window lengths) makes it hard to interpret. Figure 2 shows a case where different scales (window lengths) result in different coupling estimation, sometimes even with a different sign. Are the two signals correlated or anticorrelated and to what extent? It is not possible to draw a clear conclusion. We believe that MDC_3_ could offer a mathematically sound solution to this problem.

**Figure 2.**
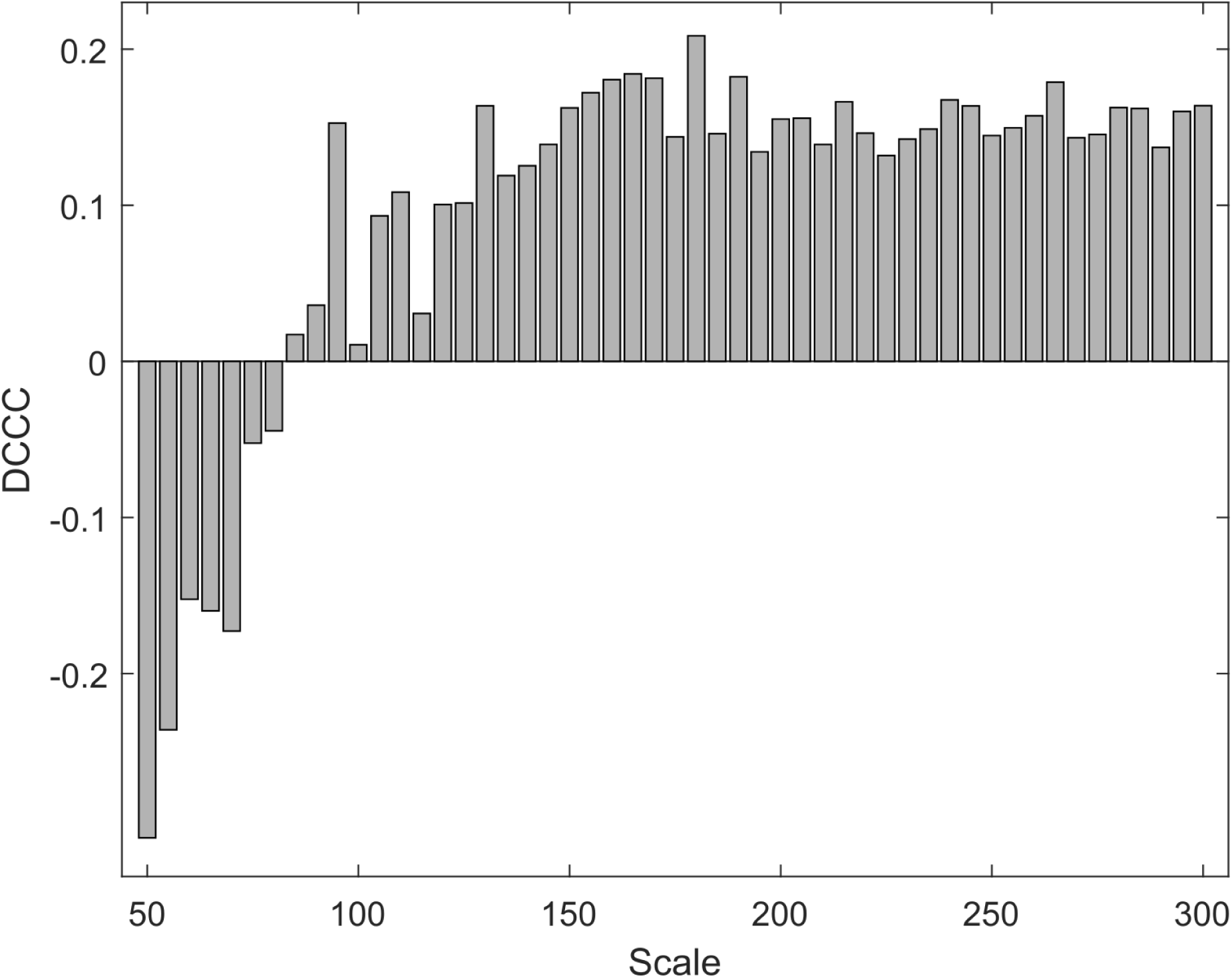
|| Detrended cross-correlation coefficient (DCCC) values for a 4 seconds-long pair of MEG signals at different scales (window lengths).

The estimation of MDC_3_ starts by calculating DCCC for different scales (window lengths). To avoid any arbitrary choice of scales (window lengths), we define frequencies (*f*) for which we would like to study the coupling of the time series. These frequencies can be converted to scales (window lengths) using the sampling rate (*SR*) of the signals (*s*=*SR*/*f*). First the DCCC for every frequency is calculated. Then, the two signals are detrended – in this case as a whole – and their cross-spectral density is estimated. We finally calculate the weighted average of DCCC, based on the relative power of each frequency in the cross-spectral density.

The distribution of DCCC – similarly to *r*_P_‘s distribution – can be skewed, so DCCC values are normalized using Fisher’s *z* transform (21,22) before the calculation of the weighted average. Details about MDC_3_ can be found in Figure 3 and the pseudo-code in **Table 1**. In this form MDC_3_ cannot construct directed graphs, i.e. the FC matrix obtained is symmetric. Using cross-covariance we can extend MDC_3_ and create directed graphs. Details about this directed variant can be found in the **Appendix**. MATLAB, Python, and R versions of MDC_3_ are available at: https://github.com/BrainModes/mdc3 (The code will be made available upon the acceptance of the manuscript).

**Figure 3.**
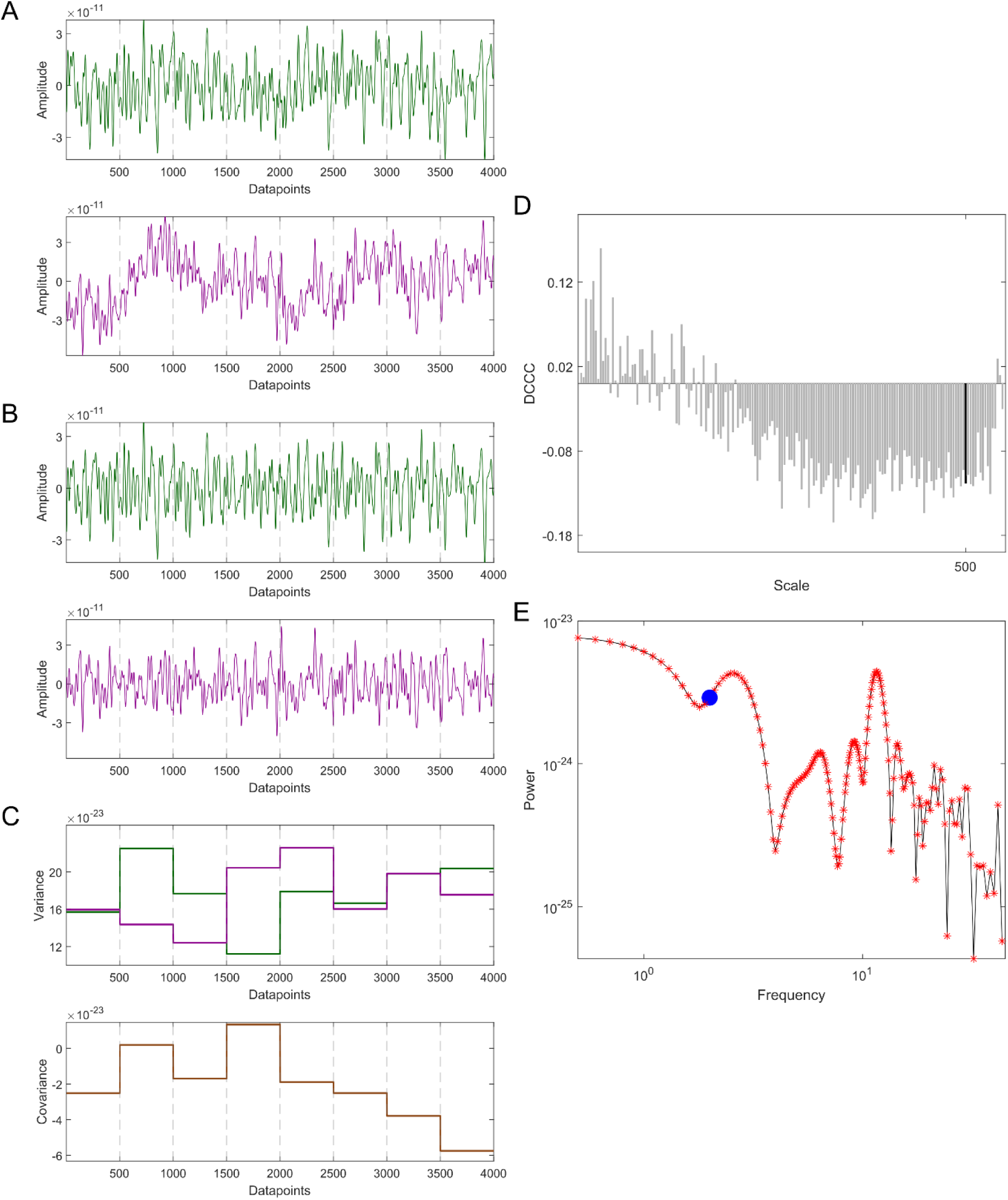
|| Demonstration of multiscale detrended cross-correlation coefficient (MDC3) using a 4 seconds-long pair of MEG signals with a sampling rate of 1000 Hz**. A**: The two signals (green and purple) are divided into smaller non-overlapping windows of length *s*, in this example *s*=500. **B**: Each window is detrended. **C**: The variances (upper panel) and covariance (lower panel) are calculated for every window. **D**: The detrended cross-correlation coefficient (DCCC) is estimated for several scales (window lengths). The black bar is the DCCC when *s*=500. **E**: The cross-spectral density of the two time series is calculated. The red asterisks correspond to the frequencies used for the estimation of DCCC, while the blue disk corresponds to 2Hz (i.e., *s*=500). MDC3 is calculated by taking the weighted average of DCCC, where the weight of each frequency is defined by the relative proportion of its power to the total cross-spectral power.

**Table 1.** || Multiscale detrended cross-correlation coefficient (MDC_3_) pseudo-code

|  |
| --- |
| **INPUTS:** time series X; time series Y; minimum frequency; maximum frequency; frequency step; sampling rate; detrending degree<br>frequencies = ([minimum frequency, maximum frequency], increment = frequency step)<br>scales = sampling rate / frequencies |
| **for** every window length |
| **for** every non-overlapping window<br>detrend (window of time series X, window of time series Y, degree = detrending degree)<br>covariance XY (window of time series X, window of time series Y)<br>variance X (window of time series X )<br>variance Y (window of time series Y) |
| $dccc = \text{mean}(\text{covariance XY}) / \sqrt{[\text{mean}(\text{variance X}) * \text{mean}(\text{variance Y})]}$ |
| [detrended X, detrended Y] = detrend (time series X, time series Y, degree = detrending degree)<br>power of frequencies = cross-spectral density (detrended X, detrended Y)<br>weights = power of each frequency / sum(power of frequencies)<br>$MDC_3 = \tanh \{ \sum [\tanh^{-1}(dccc) * \text{weights}] \}$<br>**OUTPUT:** $MDC_3$ |

### Simulated Time Series

#### ARFIMA Processes

In order to validate the efficacy of MDC_3_ we simulated pairs of auto-regressive fractionally integrated moving-average (ARFIMA) processes with known cross-correlation, as in Kristoufek (20). These series are created as follows:

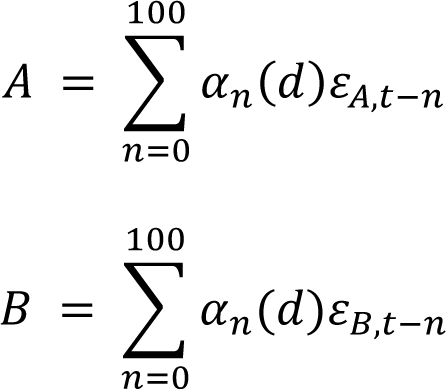

𝜀_𝐴_ is sampled from a standard normal distribution. In order to inject cross-correlation (see **Appendix** for proof) *ρ* between the two time series, we set 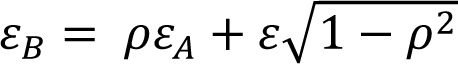, with 𝜀 being sampled from a standard normal distribution. 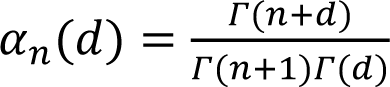, where Γ is the gamma 𝛤(𝑛+1)𝛤(𝑑) function. The parameter 𝑑 defines the non-stationarity of the simulated signal; 𝑑 < 0.5 corresponds to stationary time series, 𝑑 ≥ 0.5 corresponds to non-stationary time series. Higher values of 𝑑 indicate a higher level of non-stationarity.

We wanted to study the coupling for both stationary and non-stationary time series. So we employed the same parameters as Kristoufek (20): *i*) 𝑑 = [0.1,1.4] with increments of 0.1 and *ii*) 𝜌 = [−0.9,0.9] with increments of 0.1. To demonstrate the benefits of MDC_3_ in real-life neuronal time series, our simulations consisted of two types. The *first type* aimed to emulate EEG/MEG signals with three different lengths: 1000, 5000 & 10000 data points. We assumed that their sampling rate was 250 Hz, corresponding to 4, 20 & 40 seconds of recordings. MDC_3_ was calculated for frequencies between 0.5 and 31 Hz with increments of 0.5. In the *second type,* we wanted to study how lower sampling rates, seen in fMRI, will affect our methodology. The created signals consisted of 100, 200 & 500 data points. In this case we assumed that the sampling rate was 1Hz, meaning that the simulated time series corresponded to 100, 200 and & 500 seconds. MDC_3_ was calculated for frequencies between 0.01 to 0.12 Hz with increments of 0.01. In both types, the maximum frequencies were selected so there were at least 8 data points in every window. We decided to detrend the time series using a second-degree polynomial, since preliminary analysis showed better results compared to linear detrending. We ran 1000 simulations for each model.

We wanted to see how closely the two estimators (MDC_3_ and *r*_P_) are to the real coupling. For every 𝑑, *ρ* and signal length we calculated the root mean squared error (RMSE) of MDC_3_ and *r*_P_. Then, simulations of the same 𝑑 and signal length were grouped together. As a result, we ended up with 14 pairs (one for each value of d) of 19-points (one for each value of ρ) distributions, for every signal length (see Figure 4 for a graphical representation of the distributions). We compared every pair of distributions using a paired t-test or Wilcoxon signed rank test, depending on the normality of the underlying distributions (evaluated using Lilliefors test). Finally, Benjamini-Hochberg (BH) correction (23) was used to counteract the effect of multiple comparisons. Throughout the manuscript a comparison was considered statistically significant when BH-adjusted *p*<0.05.

**Figure 4.**
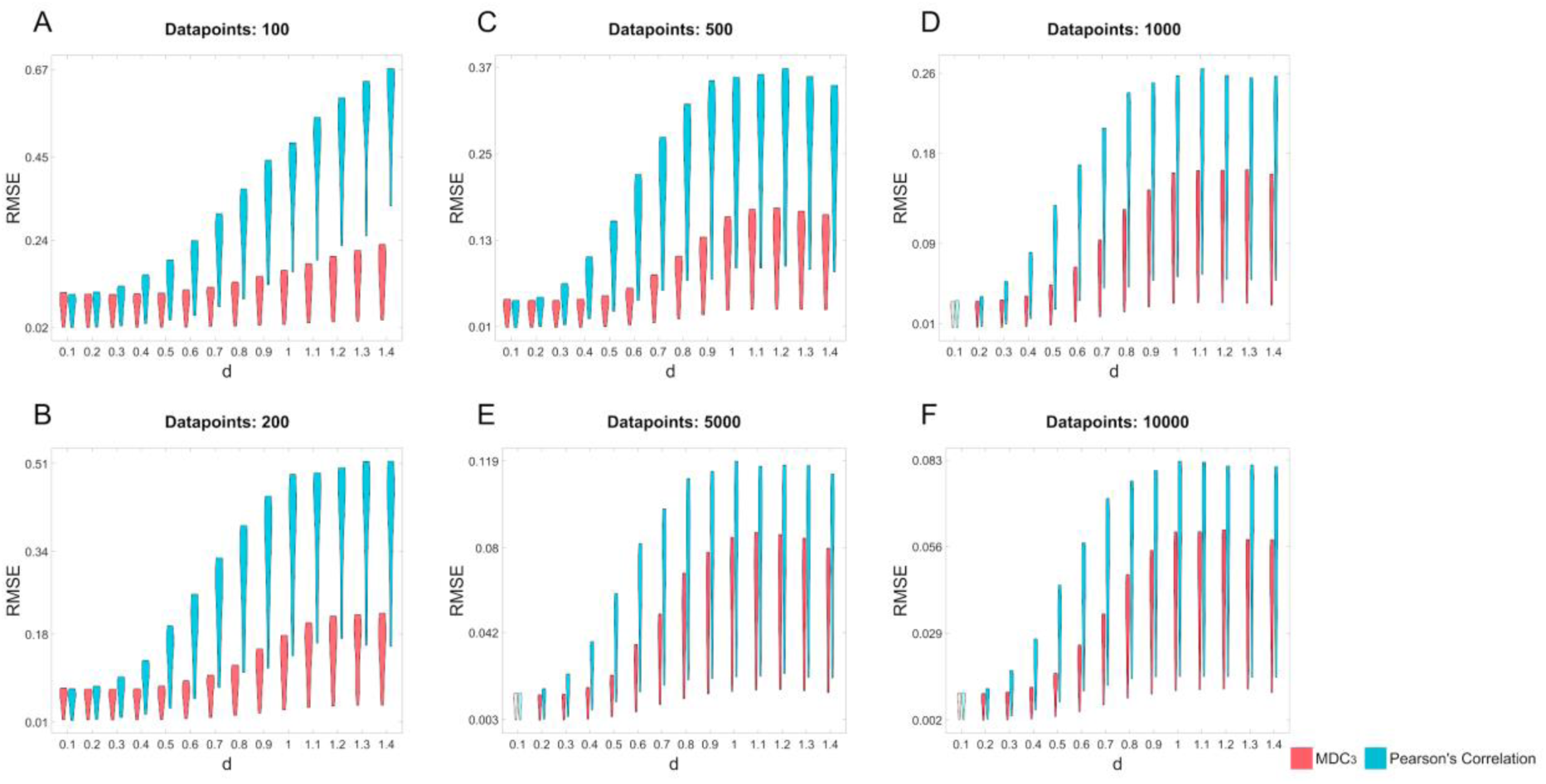
|| *Root mean squared error (RMSE) of MDC3 and Pearson’s correlation for different levels of non-stationairity (d) and signal length (panels A-F).* We simulated auto-regressive fractionally integrated moving-average (ARFIMA) processes with varying *d*, signal length and coupling strength (ρ). ρ was used to estimate the RMSE of MDC3 and Pearson’s correlation. Pairs of distributions whose difference was statistically significant (Benjamini-Hochberg adjusted *p*<0.05) are fully colored.

#### Simulated fMRI

While ARFIMA processes can create signals with known coupling, they do not represent realistic neuronal time series. For this reason, we decided to estimate the EC of fMRI signals and contrast it with the directed variant of MDC_3_. One of the most widely used EC estimators is dynamic causal modeling (DCM) (24), which estimates EC based on the constraints set by a SC matrix. Acquisition of both SC matrices (through diffusion tensor imaging) and fMRI is a lengthy and costly procedure. Thankfully, recent developments in the field of brain simulation speed up this process. We simulated the fMRI of 100 “subjects” using The Virtual Brain (TVB) (25,26). Based on the SC matrix of each subject (see next paragraph), we simulated the fMRI signal of 68 brain regions – according to the Desikan-Killiany atlas (27) – using the Reduced Wong Wang (28) neural mass model:

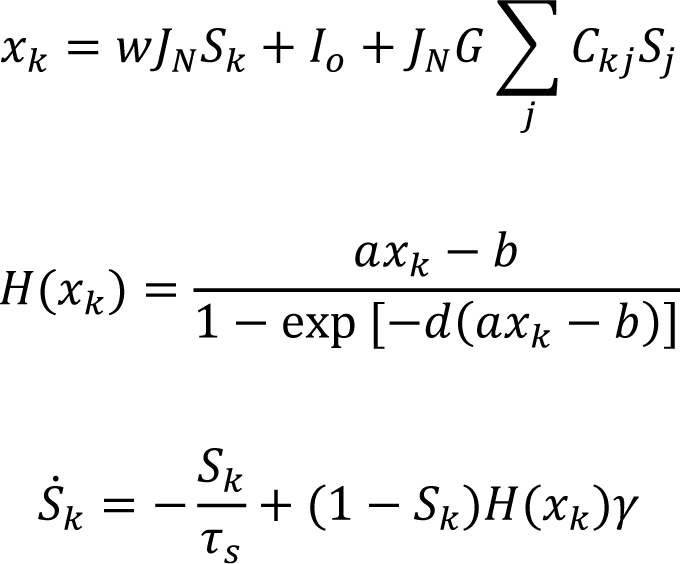

𝐻(𝑥_𝑘_) and 𝑆_𝑘_ correspond to the firing rate and synaptic gating variable of the population at the k^th^ cerebral region, respectively. 𝐺 is a global scaling factor and 𝐶_𝑘𝑗_ is the structural connection strength between the k^th^ and j^th^ regions. The description and default values of the rest of parameters can be found in Table 12 of Sanz-Leon et al. (29).

The simulated SC matrices were based the real SC matrix retrieved from https://zenodo.org/record/4263723#.Y7-8Q-zMLMI (found in “QL_20120814_Connectivity.zip”). The real SC matrix was divided into 4 quadrants. The values within each quadrant were randomly shuffled. Additionally, 30% of the connections of each quadrant were changed. Their new value was randomly selected from a normal distribution of mean and standard deviation based on the SC values of each quadrant. This shuffling and random allocation of values was also done in the accompanying tract lengths matrix created after loading “QL_20120814_Connectivity.zip” on TVB. These steps ensured that the simulated brains were different enough from the template, bust they were still biologically plausible. We then proceeded with simulating 21 minutes of fMRI time series using the Reduced Wong Wang model. The selection of appropriate parameters in brain simulations is crucial. A common practice is to perform a grid search with different combinations of parameters and compare it to properties of empirical brain activity. We varied G, w and J, while using the default values of the rest of the parameters. G was in the [0.1,29.9] range with increments of 0.1. J was in the [0,1] range with increments of 0.1. Finally, J was in the [0.2609, 0.4609] range with increments of 0.05. We estimated the FC matrix of each simulated fMRI dataset using *r*_P_. We also estimated the FC of the empirical fMRI signal (also retrieved from https://zenodo.org/record/4263723#.Y7-8Q-zMLMI) using *r*_P_. We then compared the similarities of empirical and simulated FC using Spearman’s correlation. The most realistic simulation (Spearman’s correlation 0.34) was produced when G=0.2, w=0.1 and J=0.42 while the rest of the parameters were kept in their default values.

After obtaining the simulated fMRI signals, we could proceed with the comparison between MDC_3_ and *r*_P_. While FC is simple to understand and estimate, it is merely a statistical relationship between signals. On the other hand, DCM’s constraints allow for a depiction of brain connectivity based on a more detailed network model of the brain. Hence, the EC – as captured by DCM –was chosen as the ground truth of our comparison. In DCM a realistic SC connectivity matrix is used as a template. Applying a forward model to the underlying SC can simulate fMRI signals. A parameter of this forward model is an EC matrix, which can be fine-tuned in order to produce realistic fMRI time series. Investigation of whole-brain networks with traditional DCM is a time-consuming process, which can be accelerated with regression dynamic causal modeling (rDCM) (30–32) [available at the Translational Algorithms for Psychiatry-Advancing Science (TAPAS) toolbox (33)]. rDCM offers a simplified version of DCM without severe loss in accuracy [for further details please see Frässle et al.]. In order to study the effect of signal length we analyzed the first 5, 10, 15 and 20 minutes of the simulated fMRI. This resulted in 12 matrices (4 signal lengths x 3 metrics) (**Table 2**) for every simulated brain. Since the EC matrix is not constrained between -1 and 1 as MDC_3_ and *r*_P_, we calculated the Z-scores of every EC, MDC_3_ and *r*_P_ matrix, which we then used for the comparisons. Using EC as our ground truth, we calculated the RMSE of MDC_3_ and *r*_P_ for each simulation. This resulted in 8 (2 FC estimators x 4 signal lengths)100-point (100 simulated brains) distributions. We compared every pair of distributions using a paired t-test or Wilcoxon signed rank test, depending on the normality of the underlying distributions (evaluated using Lilliefors test). The 4 *p* values were adjusted using BH correction. MDC_3_ was calculated for the frequencies between 0.011 to 0.17 Hz with increments of 0.01. 0.17 Hz was selected as the highest cutoff so each window during the estimation of MDC_3_ had 8 datapoints. Second-degree polynomials were fitted for the detrending in MDC3.

**Table 2.** *|| Demonstration of the connectivity matrices used in the analysis of simulated fMRI signals.* Multiscale detrended cross-correlation (MDC_3_), Pearson’s correlation (*r*_P_) and regression dynamic causal modeling (rDCM) were used to obtain connectivity matrices of the simulated fMRI signals. In every subject, the matrices were obtained for the first 5, 10, 15 and 20 minutes (Min) of the signal.

|  |  |  |  |
| --- | --- | --- | --- |
| 5 Min $MDC_3$ | 10 Min $MDC_3$ | 15 Min $MDC_3$ | 20 Min $MDC_3$ |
| 5 Min rDCM | 10 Min rDCM | 15 Min rDCM | 20 Min rDCM |
| 5 Min $r_P$ | 10 Min $r_P$ | 15 Min $r_P$ | 20 Min $r_P$ |

### Empirical Time Series

#### MEG Dataset

The MEG dataset consisted of eyes closed resting-state recordings of 20 elderly healthy participants (12 females, aged 71.5 ± 4.03 years), acquired using a 306-channel (102 magnetometers and 204 planar gradiometers) Vectorview MEG system (Elekta AB, Stockholm, Sweden) placed inside a magnetically shielded room (VacuumSchmelze GmbH, Hanau, Germany) located at the Laboratory of Cognitive and Computational Neuroscience (Madrid, Spain). MEG data were acquired with a sampling rate of 1000 Hz and an online [0.1 - 330] Hz anti-alias band-pass filter. All participants provided informed consent. To allow subject-specific source reconstruction, individual T1-weighted MRI scans were also available for each participant. MRI images were recorded at the Hospital Universitario Clínico San Carlos (Madrid, Spain) using a 1.5 T General Electric MRI scanner with a high-resolution antenna and a homogenization PURE filter (fast spoiled gradient echo sequence, with parameters: repetition time/echo time/inversion time = 11.2/4.2/450 ms; flip angle = 12°; slice thickness = 1 mm; 256×256 matrix; and field of view = 256 mm).

The MEG recordings were preprocessed offline using a tempo-spatial filtering algorithm (Taulu and Hari 2009) (Maxfilter Software v2.2, correlation limit of 0.9 and correlation window of 10 s) to eliminate magnetic noises and compensate for head movements during the recording. The continuous MEG data were imported into MATLAB (R2017b, Mathworks, Inc.) using the Fieldtrip Toolbox (34) (https://www.fieldtriptoolbox.org/). An independent component-based algorithm was used to remove the effects of ocular and cardiac signals from the data, together with external noises. Source reconstruction was performed using minimum norm estimates (35) with the software Brainstorm (36) (https://neuroimage.usc.edu/brainstorm/). In order to model the orientation of macrocolumns of pyramidal neurons the dipole orientations were considered to be normal to the cortical surface of the participant [see (37)]. Neural time series were finally collapsed to the regions of interest (ROI) of the Desikan-Killiany atlas (27). The data were band-pass filtered between 0.5 and 45 Hz using FIR filtering.

For every participant we analyzed multiple (ranging from 45 to 61) 4 seconds segments. We estimated the FC of each segment using MDC_3_ and *r*_P_. Then, we calculated the node strength of the brain regions by summing up the strength of every incoming and outgoing connection for every cortical area. Finally, we averaged the node strengths for all segments, so every participant had one set of node strength values. Again, we employed a series of paired t-tests or Wilcoxon signed rank tests – depending on the normality of the distributions (Lilliefors test) – to compare the node strengths of the MDC_3_ and *r*_P_ created networks. The *p*-values of each comparison group were adjusted using BH correction. MDC3 was calculated for the frequencies between 0.5 and 45 Hz. Second-degree polynomials were fitted for the detrending in MDC_3_.

#### fMRI Dataset

Finally, we analyzed 767 healthy, young adults (426 females) from the Human Connectome Project (HCP) (38). The fMRI time series were already preprocessed according to the HCP standards (39). Details about the participants can be found in the attached CSV file in the **Supplementary Information (fMRI Subjects Information)**.

For the FC estimation we used only the first eyes open resting-state period of 14.4 minutes. The dataset had a left-to-right and right-to-left echo-planar imaging (EPI) encoding. We calculated the FC using MDC_3_ and *r*_P_ for both EPI. We then averaged the FC matrices of the two EPI using Fisher’s *z* transform, as suggested by Smith et al. (38). This resulted in one MDC_3_ and one *r*_P_ FC matrix per subject. We compared the strength of each connection through a series of Wilcoxon signed rank tests that were later corrected using BH. MDC_3_ was calculated for the frequencies between 0.011 to 0.17 Hz with increments of 0.01. 0.17 Hz was selected as the highest cutoff, so each window had 8 datapoints. Second degree polynomials were fitted for the detrending in MDC3.

## Results

### Simulated Time Series

As shown in Figure 4 MDC_3_ is a more accurate estimator of coupling in the simulated ARFIMA signals in almost every case. Only some small difference can be observed for stationary signals (*d* <0.5); but as we transition to non-stationary time series (*d* ≥0.5), the RMSE of *r*_P_ is significantly higher.

The same results can be seen in realistic fMRI simulations. As Figure 5 shows, the RMSE was significantly smaller when MDC_3_ was used as an FC estimator in all signal lengths. We also see that as the signal length increases, the RMSE of *r*_P_ increases while the RMSE of MDC_3_ decreases.

**Figure 5.**
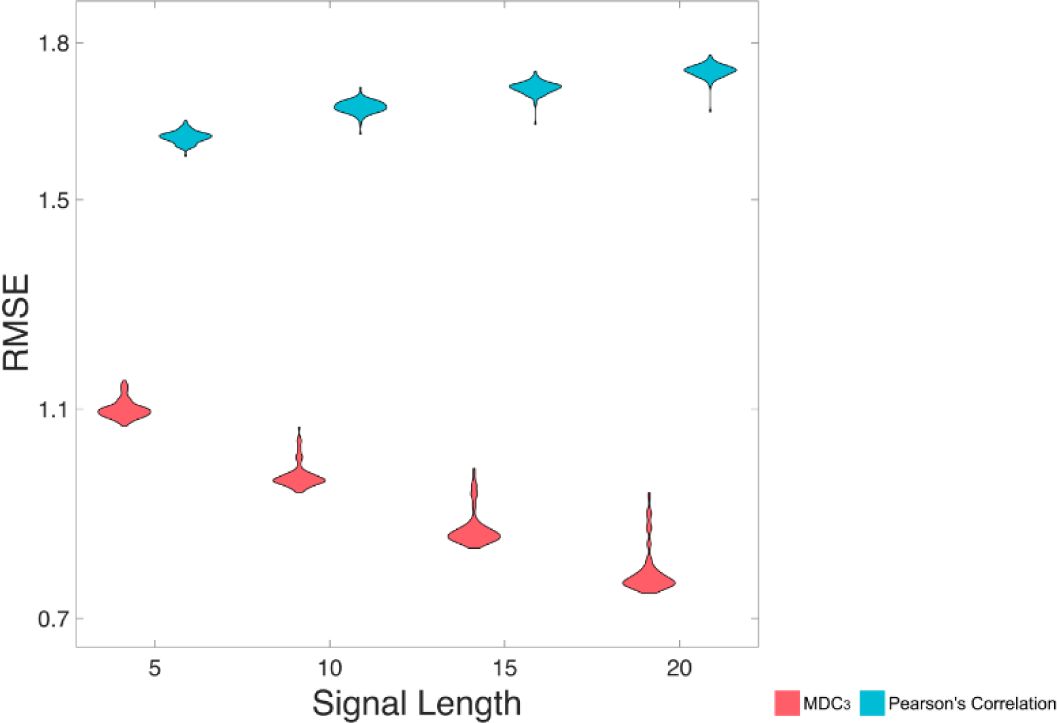
|| *Root mean squared error (RMSE) of multiscale detrended cross-correlation coefficient (MDC3) and Pearson’s correlation, for four different signal lengths (5 minutes, 10 minutes, 15 minutes and 20 minutes).* We simulated realistic fMRI signals using The Virtual Brain. The effective connectivity of the simulated brains – calculated using regression dynamic causal modeling (rDCM)– was used to estimate the RMSE of MDC3 and Pearson’s correlation.

### Neurophysiological Time Series

Figure 6 shows the difference of the node strengths between the MDC_3_ and *r*_P_ networks as estimated using MEG tracings. Significant differences can be seen in 7 channels (10%), where the *r*_P_ network had mainly higher node strengths seen by the blue color.

**Figure 6.**
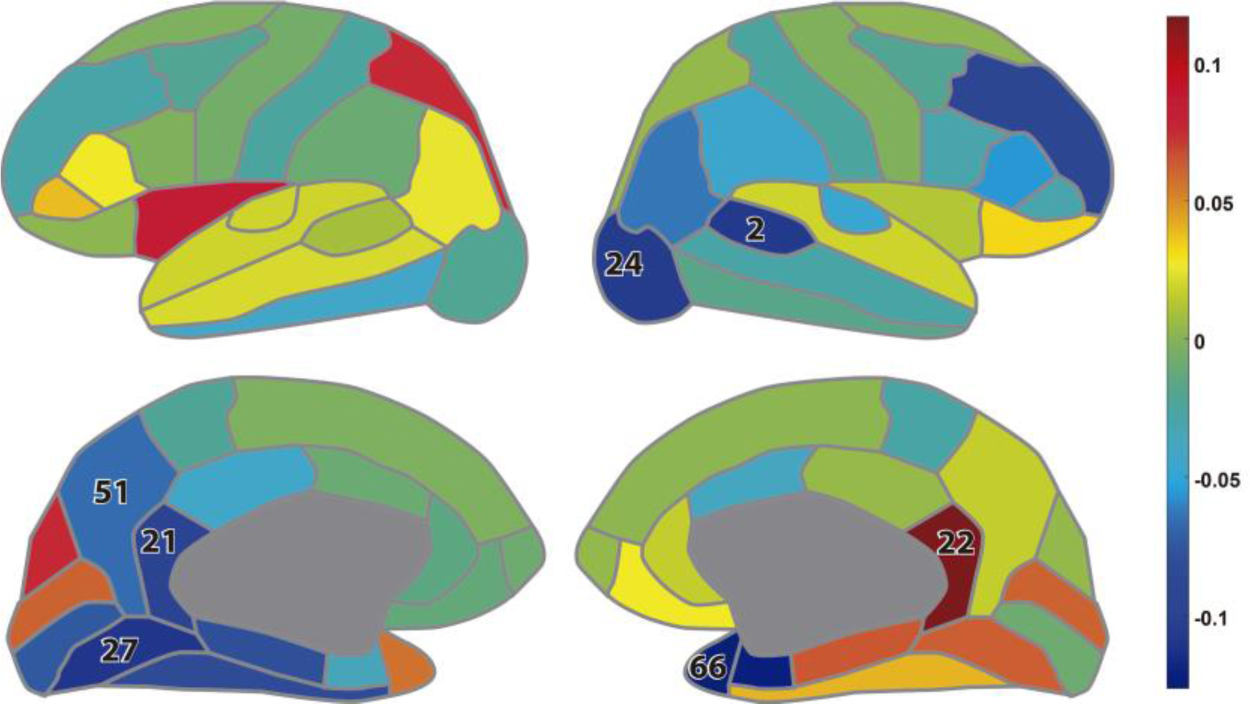
|| *Difference between the node strengths calculated during eyes closed resting-state magnetoencephalography: lateral view (up); medial view (down).* The colors represent the difference (MDC3- rP) in the node strengths while the numbers indicate the brain regions whose node strength was significantly different between the two estimators (BH-adjusted p*<0.05*). The numbers correspond to the regions of interest as defined in the Desikan-Killiany atlas (27), list provided in the **Supplementary Information (Additional Analysis)**.

For the last real-life dataset, we analyzed fMRI recordings from HCP. As Figure 7 shows, the two networks had different connectivity strengths. In some instances, *r*_P_ found higher coupling than MDC_3_ and in some other cases lower. These observations were validated statistically, since 97% (69599 out of 71631) of the comparisons were significantly different.

**Figure 7.**
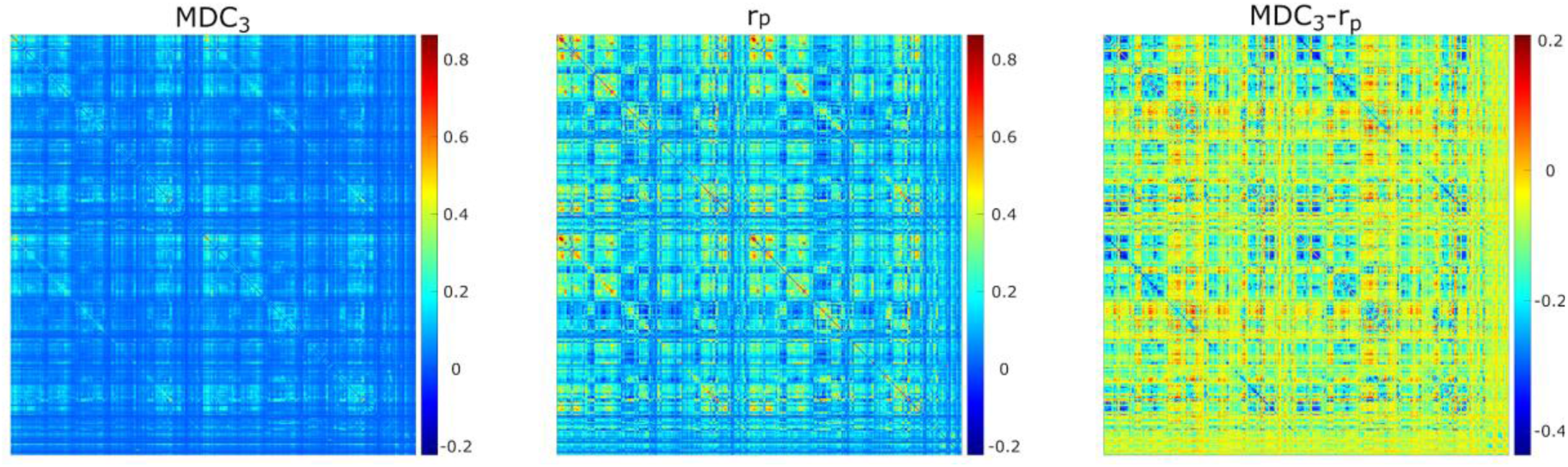
|| Averaged functional connectivity matrices using multiscale detrended-cross correlation coefficient (MDC3), Pearson’s correlation (rP), and the difference between them (MDC3-rP) using eyes open resting-state functional magnetic resonance imaging.

## Discussion

In this study we introduced the statistical metric MDC_3_ – a weighted average of DCCC – for estimating coupling in a system. Our simulations with signals of known coupling showed that MDC_3_ is a more accurate estimator of the model’s coupling parameters than *r*_P_. The exemplary FC analysis of MEG and fMRI data also showed that the use of MDC_3_ could lead to significant differences in the connectivity matrices compared to *r*_P_.

We simulated 1000 pairs of time series of different coupling strengths, signal lengths and degrees of non-stationarity. For each pair we calculated MDC_3_ and *r*_P_. As explained in the **Introduction**, and shown in Figure 1, the variance and covariance of stationary signals remain constant, meaning that MDC_3_ and *r*_P_ will be similar. This is not the case for non-stationary series whose variance and covariance heavily depend on time. Our simulations confirm that, since the RMSE of MDC_3_ was significantly smaller in every case, except for fairly stationary signals (Figure 4). The discrepancy between the two estimators increased greatly with higher levels of non-stationarity. Similar findings have been reported for DCCC in Kristoufek (20). We also simulated a series of fMRI signals using TVB. We could not simulate realistic neuronal time series with known coupling, so we decided to use the EC matrices of the simulations as ground truth. The results showed that MDC_3_ is closer to the EC compared to *r*_P_ (Figure 5). We also observed that as the length of the signals increased the accuracy of MDC_3_ increased, contrary to *r*_P_. Smith et al. (40) decided to validate FC estimators using the underlying SC as ground truth. While we considered this approach, we decided to use EC instead. The choice was based on the two following reasons. Firstly, SC cannot entirely predict FC (41). Secondly, the lack of negative values in SC would not allow for accurate study of negatively correlated brain regions. For the sake of completeness, we also compared MDC3 and *r*P of the simulated fMRI signals using SC as ground truth. This time, *r*P was found to be a better estimator, albeit with a narrow margin (see **Additional Analysis** in **Supplementary Information**). An interesting byproduct of this analysis was that *r*_P_ was similar to SC, while EC and MDC_3_ were similar to the tract length matrices used for the construction of the simulations. While this finding is interesting, it is beyond the scope of this study and should be revised in future studies. The matrices of each simulation can be found in the **Supplementary Information** (**TVBMatrices**). Finally, we repeated our MDC_3_ and *r*_P_ comparisons this time using the simulations from Smith et al. (40). In the majority of cases MDC_3_ was more accurate, especially when EC was used as ground truth. The complete results of the additional analysis can be found in the **Supplementary Information (Additional Analysis).**

Of course, statistical significance in simulations without real-life benefits would not warrant the use of MDC_3_. To demonstrate its advantages, we used MEG and fMRI datasets. As shown in Figure 6, using MDC_3_ and *r*_P_ as FC estimators resulted in significantly different brain networks. In some cases, the node strengths of the *r*_P_ networks were higher, while in others they were lower. After analyzing the FC matrices of the fMRI dataset, we saw that almost all connections were significantly different between the two matrices (Figure 7). Once again, some connections were stronger and some weaker when *r*_P_ was used. A homogenous overestimation or underestimation would not have been a serious drawback since FC studies usually rely on relative comparisons and not on the exact values themselves. But it seems that in some regions *r*_P_ would give lower values and in others higher, presenting a rather false picture of the brain network. At a first glance, someone might be dismissive of this, since it is well known that different estimators can lead to different FC matrices (11–13). This would have been the case if we had not seen the higher reliability of MDC_3_ both from a mathematical standpoint (**Methods**) and in simulations (**Results**). We then suggest that MDC_3_ should be preferred over *r*_P_. Even if MDC_3_ is computationally more expensive, today’s computational capabilities make the time difference negligible.

Finally, it should be noted that MDC_3_ is still a linear FC estimator. Non-linear estimators like PLV, MI, PLI, and SL still capture dynamics that MDC_3_ cannot. In spite of that, we believe that MDC_3_ is a valuable addition to the FC field due to its ability to capture the sign of correlation (i.e., correlation vs anticorrelation); something that the aforementioned non-linear estimators cannot do. A common practice in FC studies is the exclusion of anticorrelations (4). Since the human brain operates with several negative feedback loops, we believe that it is necessary to study anticorrelation in order to obtain more accurate brain architectures, as suggested by previous studies (42,43). We decided to explore this further in the **Supplementary Information (Additional Analysis)** using the MEG dataset. Briefly, we compared the FC matrices as estimated with MDC_3_ and PLV using two different source reconstruction pipelines, i.e., with constrained and unconstrained dipoles. The first method makes it possible to obtain a more realistic phase (and sign) of the reconstructed time series. This benefit can be overshadowed by the inability of most FC estimators to capture the sign of coupling, including PLV. As a result, such metrics could mistakenly identify correlation for anticorrelation and vice versa. As expected, MDC_3_ detected more differences between the reconstructions with constrained and unconstrained dipoles than PLV.

DCCC and its extension MDC_3_ are closely related to the scale-free analysis of signals. The numerator and denominator of **Equation 1** are integral parts of the detrended fluctuation analysis (44) and detrended cross-correlation analysis (45) analysis, respectively. DCCC has been incorporated in surrogate testing of fractal (scale-free) coupling already (12,19,46–48). The main difference between the two methods is the single output of MDC_3_, as opposed to scale-specific correlations of DCCC. It is then clear that MDC_3_ cannot be used in surrogate testing of fractal FC, since scale-specific estimators are necessary for such analysis. DCCC has also been employed in multifractal FC (49); where different exponents capture different sizes of fluctuations. Theoretically, a multifractal MDC_3_ could be created as well. This is beyond the scope of the current study because we focused on improving the interpretability of DCCC. The calculation of MDC_3_ using different scaling exponents would add another layer of complexity to the interpretation of the outputs. Recently, a real-time algorithm for the estimation of DCCC was presented (50,51), which can be extended for MDC_3_ as well. This means that MDC_3_ can be used in brain-computer interfaces or clinical monitoring of patients, where constant tracking of network dynamics is needed.

## Conclusion

We presented a new estimator of coupling between time series termed multiscale detrended cross-correlation coefficient. Using simulated data, we showed a higher accuracy over *r*_P_. The differences between the two estimators were made apparent in MEG and fMRI datasets of healthy populations. Here we explored the benefits of MDC_3_ only in neuronal time series. We believe that our new method has the potential to be used in several other disciplines where linear coupling of non-stationary signals is investigated. Of course, appropriate validation pipelines specific to each field are recommended before any prior use.

## Appendix

### Auto-Regressive Fractionally Integrated Moving-Average Processes

Assume two distributions 𝜀_𝐴_ and 𝜀_𝛣_. 𝜀_𝐴_ is a standard normal distribution, meaning E[𝜀_𝐴_] = 0 and 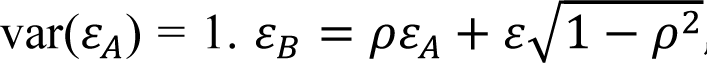, where *ε* is also standard normal [i.e. E[*ε*] = 0 and var(ε)=1]. The variance of 𝜀_𝛣_ can be calculated as follows:

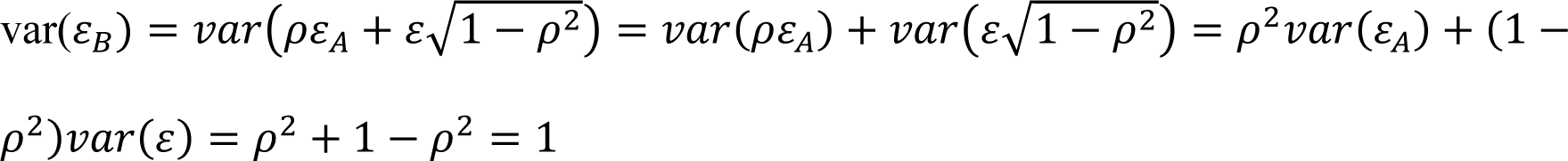

Then the real coupling between the two distributions can be calculated as:

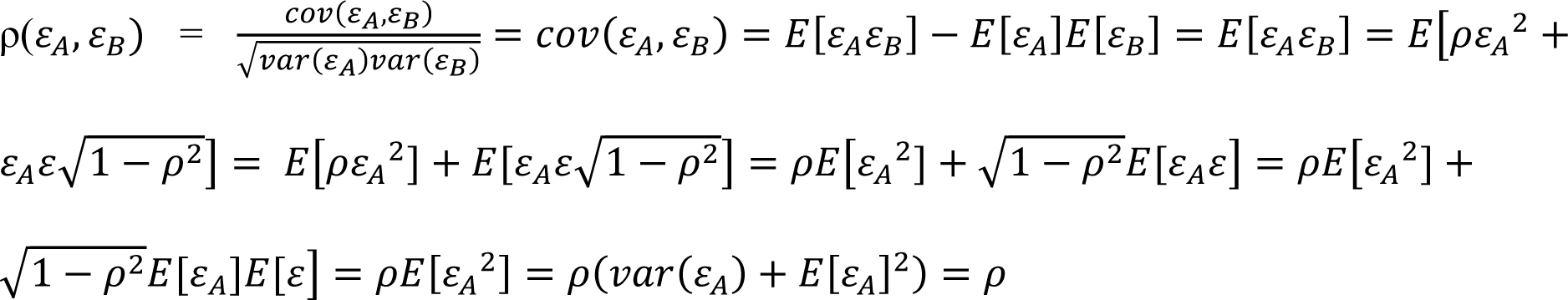

The two ARFIMA series 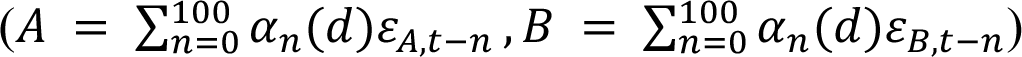 are the cumulative sums of 𝜀_𝐴_ and 𝜀_𝛣_ multiplied by a step-specific weight [(𝛼_𝑛_(𝑑)]. The only source of stochasticity of *A* and *B* are 𝜀_𝐴_and 𝜀_𝛣_, meaning that the true coupling between *A* and *B* is *ρ*.

### Directed MDC_3_

The difference of the directed variant of MDC_3_ is that for every detrended signal 𝑋̂_𝑖_ and 𝑌̂_𝑖_ the cross-covariance(𝑋̂_𝑖_, 𝑌̂_𝑖_) is estimated, instead of the covariance(𝑋̂_𝑖_, 𝑌̂_𝑖_). The maximal covariance – in absolute terms – for negative lags is used for the DCCC estimation when 𝑋̂_𝑖_ is leading. Similarly, the maximal covariance – in absolute terms – for positive lags is used for the DCCC estimation when 𝑋̂_𝑖_ is following.

## Author Contributions

O.S. developed MDC3, wrote the MATLAB, R and Python code for MDC3, performed data analysis and interpretation, and wrote the first draft of the manuscript. G.S. performed data analysis and interpretation. M.H. contributed to data interpretation. I.S.M. performed data analysis and interpretation. D.L-S. performed data analysis and interpretation M.S. performed data analysis and interpretation. P.R. provided conceptual guidance, supervision and funding throughout the study. All authors contributed to reviewing the manuscript and approved its final version.

## Financial Disclosure Statement

P.R. acknowledges support by Digital Europe TEF-Health 101100700, EU H2020 Virtual Brain Cloud 826421, Human Brain Project SGA2 785907; Human Brain Project SGA3 945539, ERC Consolidator 683049; German Research Foundation SFB 1436 (project ID 425899996); SFB 1315 (project ID 327654276); SFB 936 (project ID 178316478; SFB-TRR 295 (project ID 424778381); SPP Computational Connectomics RI 2073/6-1, RI 2073/10-2, RI 2073/9-1; PHRASE Horizon EIC grant 101058240; Berlin Institute of Health & Foundation Charité, Johanna Quandt Excellence Initiative; ERAPerMed Pattern- Cog, the Virtual Research Environment at the Charité Berlin – a node of EBRAINS Health Data Cloud.

## Notes

### Competing Interest Statement

The authors have declared no competing interest.

